# GRINtrode: A neural implant for simultaneous two-photon imaging and extracellular electrophysiology in freely moving animals

**DOI:** 10.1101/2022.06.17.496591

**Authors:** Connor M. McCullough, Daniel Ramirez-Gordillo, Michael Hall, Gregory L. Futia, Andrew K. Moran, Emily A. Gibson, Diego Restrepo

## Abstract

**Significance:** In vivo imaging and electrophysiology are powerful tools to explore neuronal function that each offer unique complementary information with advantages and limitations. Capturing both data types from the same neural population in the freely moving animal would allow researchers to take advantage of the capabilities of both modalities and further understand how they relate to each other.

**Aim:** Here we present a head-mounted neural implant suitable for in vivo two-photon imaging of neuronal activity with simultaneous extracellular electrical recording in head-fixed or freely moving animals.

**Approach:** A GRIN lens-based head-mounted neural implant with extracellular electrical recording provided by tetrodes on the periphery of the GRIN lens was chronically implanted. The design of the neural implant allows for recording from head-fixed animals, as well as freely moving animals by coupling the imaging system to a coherent imaging fiber bundle.

**Results:** We demonstrate simultaneous two-photon imaging of GCaMP and extracellular electrophysiology of neural activity in awake head-fixed, and freely moving mice. Using the collected information, we perform correlation analysis to reveal positive correlation between optical and local field potential recordings.

**Conclusion:** Simultaneously recording neural activity using both optical and electrical methods provides complementary information from each modality. Designs that can provide such bimodal recording in freely moving animals allow for the investigation of neural activity underlying a broader range of behavioral paradigms.

## 1 Introduction

In vivo recording of neural activity allows researchers to probe the function of neural circuits in action. The oldest method for this type of recording is electrophysiological recording of the extracellular field potential^1^. Implanted electrodes can record neural oscillations due to ensemble activity of large numbers of neurons known as local field potential (LFP), as well as spiking activity from single neurons when multiple recording sites are used^2,3,4^. Electrodes can be implanted to deep brain regions with limited damage to tissue and offer excellent temporal resolution with sampling rates in the tens of kilohertz capable of recording the changes in field potential elicited by action potential firing and synaptic activity. However, in vivo recording of LFPs has limitations in the ability to resolve single neuron activity. The recording field of an electrode is spatially limited by an inverse-square law limiting the number of neurons that can be sampled per electrode, and electrical recording does not provide high resolution spatial information regarding the position of recorded cells relative to the recording site.

Optical techniques can overcome these spatial hurdles by allowing for in vivo neural recordings at single neuron resolution. These techniques use fluorescent dyes or genetically encoded indicators with fluorescence intensity that changes as a function of intracellular ionic concentration (typically calcium)^5^, neurotransmitter release^6^, or membrane voltage^7,8^. Optical methods allow for recording from large numbers of neurons, with known spatial localization allowing researchers to study spatial network dynamics and record from the same neurons longitudinally. Expression of these fluorescent reporters can be localized to specific cell types by genetically targeted expression. However, the temporal resolution of optical recordings is limited by the dynamics of the fluorescent reporter and acquisition rate of the microscope used, and fluorescent reporters based on ionic concentration are only a proxy of spiking. In addition, optical indicators do not provide the information on coordinated neuronal activity relevant to the flow of neural information^9^ that is provided by LFP recordings.

Efforts have been made to combine optical and electrical recording modalities using transparent electrodes underlying a cranial window, but this limits the recording area to superficial structures^10^. The popular Miniscope^11^ platform has also been used to achieve simultaneous optical and electrical recordings^12^, but this method is limited to 1 photon imaging, and fabricating the device may prove difficult to labs without specialized microelectromechanical equipment for manufacturing the electrode array. More easily constructed single electrode based designs have been developed^13^, but this method is restricted to head-fixed animals and faces limitations in isolating single units presented by single electrode recording modalities. Additional methods based on silicon probes have been developed^13^ that provide multisite recording, but present a complex multi-component surgery and are limited to head-fixed mice.

Here we describe the development of a low-cost GRIN lens and tetrode-based device, that we name the GRINtrode. The GRINtrode allows for simultaneous two-photon imaging of neuronal activity using GCaMP fluorescent calcium indicators and extracellular field potential recorded from tetrodes surrounding a chronically implanted GRIN lens. The GRINtrode allows for multiphoton imaging and electrical recording in head-fixed animals, as well as in freely moving animals by coupling the imaging system to a coherent imaging fiber bundle. Tetrodes provide the advantage of multisite recording, allowing for isolation of single unit electrical activity. The GRINtrode also includes a ventral drive mechanism to advance the implant depth post-surgery. We provide examples of this dual recording and an analysis of the correlations between the fluorescent GCaMP activity traces and local field potential or single unit electrical activity.

## 2 Materials and Methods

### 2.1 GRINtrode design and fabrication

The GRINtrode components were designed in Solidworks 3D CAD software (Dassault Systèmes). Aluminum GRINtrode bodies, PEEK drive thumb screws, Delrin rods, and vented 4-40 socket head cap screws were machined by the Neuroscience Machine Shop at the University of Colorado Anschutz Medical Campus. Polyimide tubing was obtained from MicroLumen, Inc. GRIN lenses were obtained from GRINtech GmbH and Inscopix, Inc. Tetrodes are spun using a magnetic stirring plate^14^ from 12.7 μm diameter polyimide coated nichrome wire (Sandvik, PX000001), bonded using a heat gun set to 235 degrees Celsius, and impedance is set to 250-500 kΩ by gold plating using a BAK Electronics IMP-2A and Neuralynx gold plating solution. Tetrodes are fixed around the periphery of the GRIN lens in a cross pattern using Loctite Ultra Gel cyanoacrylate glue (Loctite PN 1906107) and isolated from the imaging optic channel and vented screw using concentric sections of polyimide tubing. Tetrodes were cut by hand to extend ∼200-300 μm below the implanted GRIN lens surface to place the electrical recording sites close to the imaging plane of the GRIN lens system. Electrodes were pinned to a Neuralynx EIB-16 electrode interface board and then covered with 2-part epoxy (Gorilla Glue PN 4200101).

The GRINtrode utilizes a 9 mm GRIN relay lens which is chronically implanted into the tissue with the device, referred to as the implanted GRIN lens. We have used both GRINtech 1 mm dia. x 9 mm length lenses (GRINtech NEM-100-25-10-860-S-1.0p) and Inscopix 1 mm dia. x 9 mm length lenses (Inscopix 1050-002177) for the implanted GRIN lens. Above the implanted GRIN lens is an empty polyimide tube which creates a channel to allow for an additional imaging optic to be inserted (Fig. 1A). The imaging optic channel is covered with Parafilm and sealed with Kwik-Sil silicone elastomer (WPI, order code KWIL-SIL) between imaging sessions. For head-fixed imaging, the imaging optic is another GRIN lens, referred to as the imaging GRIN lens (Figs. 1B, 1C). We have used either GRINtech 0.85 mm diameter x 10.8 mm length (NEM-085-45-10-860-S-1.5p) or GRINtech 1 mm diameter x 14.07 mm length (NEM-100-25-10-860-S-1.5p) GRIN lenses as the imaging GRIN lens.

**Figure 1.**
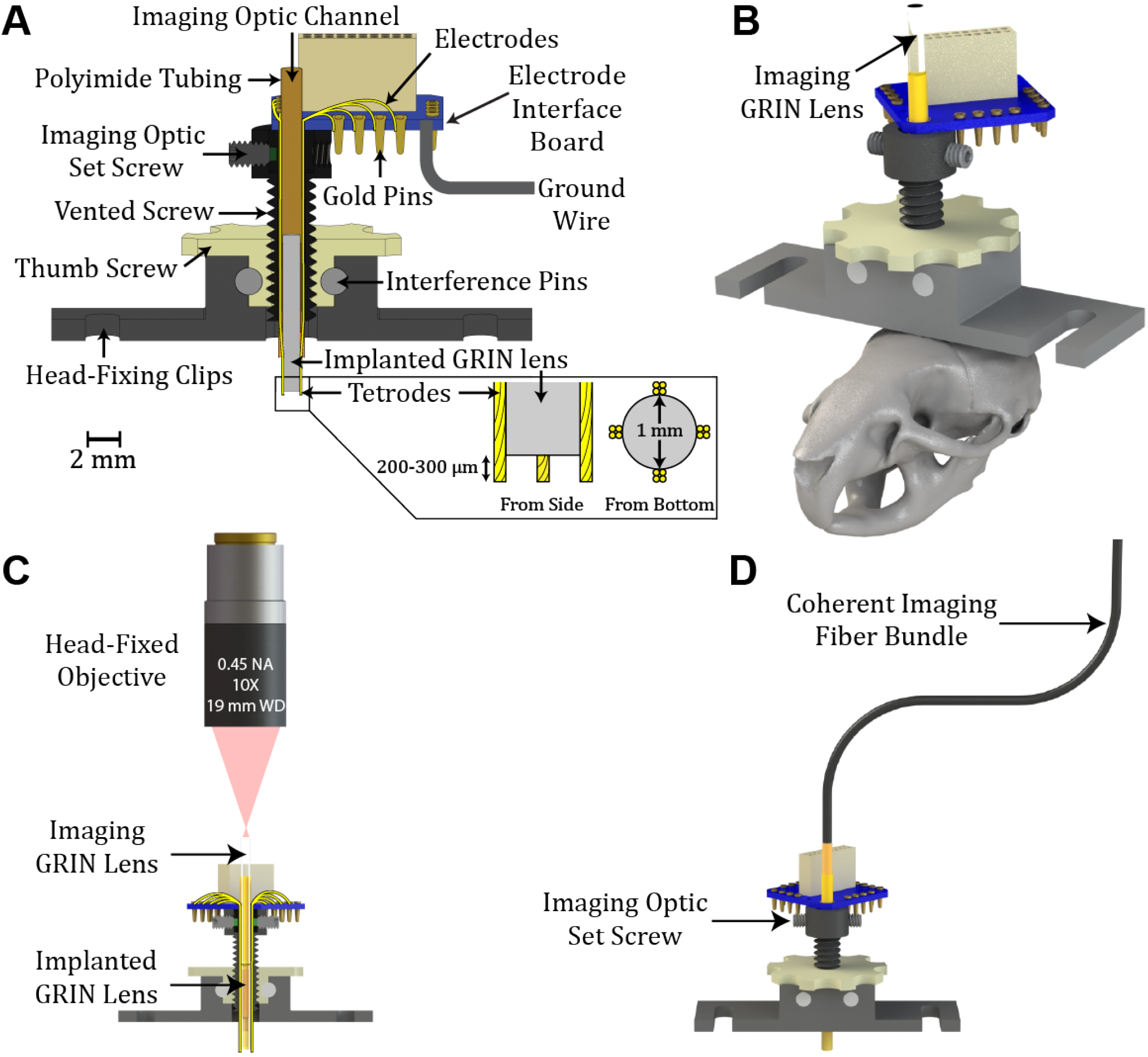
GRINtrode design. **(A)** Cross section of the GRINtrode. The GRINtrode consists of 4 major components: An aluminum main body, a PEEK drive thumb screw, a stainless steel vented 4-40 socket head cap screw, and an electrode interface board. The vented screw is threaded into the drive thumb screw, and the drive thumb screw is held in place by friction fitted Delrin interference pins. Flat sides are machined onto the vented screw, which interact with flat faces on the hole in the bottom of the aluminum body to create a vertical translation drive. Turning the drive thumb screw will increase the implantation depth of the GRINtrode probe without any rotation. The aluminum body includes an integrated head-fixing bar to improve head-fixed imaging stability and simplify the surgical procedure. Polyimide tubing is glued inside the vented screw and around GRIN lenses to provide a protected channel for tetrodes and isolate them from inserted imaging optics. The vented screw cap has threaded holes for nylon tipped #0-80 set screws that can be used to lock the imaging optic into place. **(B)** Render of the GRINtrode as it would be used for head-fixed imaging to scale on a mouse skull. The GRINtrode weighs < 3g when fully assembled and is suitable for chronic implantation with stability demonstrated for up to 6 months. **(C)** Cross-section of the head-fixed GRINtrode imaging system. A 19 mm WD objective (Edmund Optics, #58-372) is used to avoid interference with the connector attached to the electrode interface board. Additionally, a custom extension cord (Omnetics A71325-001) is used to further remove the Neuralynx RHD2132 amplifier from the imaging path. **(D)** Render of the GRINtrode in fiber coupled modality as it would be used for recording from freely moving animals. The fiber-GRIN lens spacing is adjusted using a Sutter MPC-325 micromanipulator system, and the fiber is locked into place using the imaging optic set screw.

For fiber coupled imaging, a coherent imaging fiber bundle was inserted into the imaging optic channel (Fig. 1D). A 1.5-meter-long Fujikura FIGH-15-600N coherent imaging fiber bundle was used for two-photon fiber coupled imaging. For one-photon fiber coupled imaging (Supplemental Fig. 1), the coherent imaging fiber bundle was epoxied to a GRIN doublet lens, GRINtech 1 mm dia. x 8.09 mm length doublet GRIN lens (NEM-100-25-10-860-DS). The doublet GRIN lens introduces a 2.6x magnification, which serves to demagnify the core-to-core spacing of the coherent imaging fiber bundle, improving the effective lateral resolution of the imaging system. However, using this additional GRIN doublet lens also restricts the field of view, and for two-photon fiber coupled imaging there was a decrease in power throughput. The fiber-implanted GRIN spacing was adjusted using a Sutter Instruments MPC-325 micromanipulator and the fiber was locked into place using a set screw.

### 2.2 Microscope setup

A Sutter Instruments Movable Objective Microscope was used to perform head-fixed in-vivo 2P imaging through the GRINtrode. The excitation source was a Spectra-Physics MaiTai DeepSee HP Ti:Sapphire laser with ∼80 fs pulses tuned to a center wavelength of 920 nm and operating at 80 MHz pulse repetition rate. Emission was collected through a 525/50 nm bandpass filter (Semrock PN FF03-525/50-25). For head-fixed recordings an Edmund Optics 10X High Resolution Infinity Corrected Objective (Edmund Optics, #58-372) with a 0.45 NA and 19 mm working distance was used to match the NA of GRIN lenses and provide enough working distance to avoid interference with the tetrode amplifier.

For two-photon fiber-coupled recording, we used a previously described^15^ custom-built two-photon microscope with an Olympus UPlanXApo 10X objective (Olympus, PN N5701900) focusing the imaging laser on to the surface of a 1.5 m length Fujikura FIGH-15-600N coherent imaging fiber bundle. In order to obtain short pulses at the sample, the output of the MaiTai laser was sent through a 15 cm polarization-maintaining single-mode fiber (Thorlabs PN PM780-HP) for spectral broadening and then a grating pair compressor (600 lines/mm, Newport PN 33009BK02-351R) to correct for second order dispersion. The grating compressor has been slightly modified from the previously described system, using a 15 cm polarization-maintaining single-mode fiber and 800 lines/mm gratings. The end of the distal side of the coherent imaging fiber bundle is housed in a XYZ translator (Newport OC1-TZ, Newport OC1-LH1-XY) to allow alignment of the fiber bundle surface to the focus of the microscope objective.

For one-photon fiber-coupled recording (supplemental), we used a Nikon A1R confocal microscope with an Olympus UPlanXApo 10X objective focusing the excitation laser onto a 1.5-meter-long Fujikura FIGH-15-600N coherent imaging fiber bundle. The proximal end of the Fujikura coherent imaging fiber bundle was coupled to a doublet GRIN lens (GRINtech NEM-100-25-10-860-DS) by gluing the fiber and the GRIN lens into a polyimide tube, and then inserted into the GRINtrode imaging optic channel. A XYZ translator (Thorlabs CXYZ1) allowed alignment of the fiber bundle surface to the microscope focus.

### 2.3 Tetrode recording setup

Tetrodes were passed through the GRINtrode and pinned into a Neuralynx EIB-16 electrode interface board. A custom Omnetics cable harness (Omnetics A71325-001) was connected to the EIB-16 board and then to an Intan RHD2132 16 channel amplifier (Intan C3334). The custom harness was used to extend the connection to the amplifier away from the imaging GRIN lens such that it does not interfere with the imaging path. The amplifier was then connected to an Intan SPI cable (Intan C3206) and passes to an Intan RHD USB interface board (Intan C3100). The RHD USB interface board was connected to the recording computer with a USB cable. The Intan RHD USB interface also recorded a synchronization frame trigger output from the microscope so that imaging data and electrophysiology data could be time synced.

### 2.4 Image Processing

Head-fixed image timelapses were processed with the CaImAn software package for calcium image analysis^16^. CaImAn was used to perform motion correction, identify cell ROIs, generate ΔF/F traces of GCaMP fluorescence, and reconstruct denoised timelapses. Fiber coupled 2-photon image timelapses were analyzed manually with Fiji^17^ because the fiber bundle artifact prohibited analysis with CaImAn.

### 2.5 Tetrode Data Processing

Tetrode recordings are analyzed with in-house MATLAB software (GitHub/restrepd/drta, GitHub/restrepd/drgMaster and GitHub/ConMark/GRINtrode/Code) and the Wave_clus software package^18^ modified as posted in GitHub/restrepd/wave_clus.

### 2.6 Behavioral Video Analysis

Behavioral videos of freely moving animals were recorded at 30 fps using an infrared illuminated webcam (ELP*-*USBFHD01M*-*DL36). Behavioral videos were time synced to the imaging data by isolating the frames when the infrared light from the imaging laser first and last appeared. Behavioral kinematics were extracted using the DeepLabCut^19^ markerless pose estimation software package.

### 2.7 Animals

Thy1-GCaMP6f mice^20^ (JAX RRID:IMSR_JAX:024276) were housed in a vivarium with a reversed light cycle of 14/10 h light/dark periods with lights on at 10:00 p.m. Food (Teklad Global Rodent Diet no. 2918; Harlan) was available ad libitum. All experiments were performed according to protocols approved by the University of Colorado Anschutz Medical Campus Institutional Animal Care and Use Committee.

### 2.8 Surgery

Mice 2-5 months of age were anesthetized by brief exposure to isoflurane (2.5%) and subsequently anesthesia was maintained with an intraperitoneal injection of ketamine (100 mg/kg) and xylazine (10 mg/kg). The mouse was then placed in a stereotaxic frame. A 1.6 mm diameter craniotomy was made above the target site using a dental drill. 1-2 μL of virus (pAAV1.Syn.GCaMP6s.WPRE.SV40 or pGP-AAV1.Syn.jGCaMP7f.WPRE) was injected 500 μm above the target coordinates at a rate of 100 nL per minute. After the viral injection, tissue was aspirated^21^ to ∼500 μm above the target coordinates, and the GRINtrode device was implanted at the target coordinates (see section 3.3 for coordinates) using a motorized micromanipulator (Sutter Instruments MPC-325). One ground screw was inserted 1 mm posterior from bregma and 1 mm lateral to the midline. The ground wire of the GRINtrode (A-M Systems Bare Silver Wire 0.008, Fisher Cat. No. NC0326296) was then wrapped around the ground screw and sealed with silver paint to optimize electrical contact (SPI Supplies PN 04999-AB). The ground screw and GRINtrode body were then sealed to the bone with C&B Metabond (Parkell SKU S380). Mice were allowed to recover for 4 weeks before the initiation of experiments and to allow time for virus expression.

## 3 Results

### 3.1 The GRINtrode

The GRINtrode is a head-mounted neural implant that uses a GRIN lens with surrounding tetrodes designed to provide simultaneous imaging of neuronal activity reported by GCaMP and recording of extracellular electrical neuronal activity (Fig. 1A). This is achieved by simultaneous two-photon imaging through a GRIN lens and extracellular field potential recording provided by four tetrodes placed on the periphery of the GRIN lens. The GRINtrode design is based on the optetrode^22^, with a modified housing and bracket to allow for head-fixed or multicore fiber coupled freely moving imaging and utilizing a GRIN lens instead of an optical fiber for imaging (Fig. 1B, 1C). By utilizing a GRIN lens and tetrode-based design, we provide a solution for simultaneous optical imaging and extracellular electrophysiology from deep brain structures using either head-fixed one-photon or two-photon microscopy. Additionally, the GRINtrode imaging system can be coupled to a coherent imaging fiber bundle^15^ (Fig. 1D) to allow for freely moving fiber-coupled one-photon (Supplemental Fig. S1) or two-photon microscopy (Fig. 5) using a coherent imaging fiber bundle^15^ with simultaneous extracellular electrophysiology.

### 3.2 Optical GRINtrode characterizations

Head-fixed two-photon GRINtrode imaging was performed by inserting a GRIN lens into the imaging optic channel of the GRINtrode, and imaging through two GRIN lenses in series. In the data presented, we used two different combinations of GRIN lenses. For imaging system A, the chronically implanted GRIN lens is a GRINtech 1 mm diameter, 9 mm long GRIN lens (GRINtech NEM-100-25-10-860-S-1.0p), and the imaging GRIN lens is a GRINtech 1 mm diameter, 14 mm long GRIN lens (GRINtech NEM-100-25-10-860-S-1.5p). For imaging system B, the chronically implanted GRIN lens is an Inscopix 1 mm diameter, 9 mm long GRIN lens (Inscopix 1050-002177), and the imaging GRIN lens is a GRINtech 0.85 mm diameter, 10.8 mm long GRIN lens (GRINtech NEM-085-45-10-860-S-1.5p). Imaging system B was used for an early iteration of the GRINtrode where the polyimide tubing used to create the imaging optic channel was too small in diameter to allow for 1 mm diameter imaging GRIN lenses to be inserted. In later iterations the implanted GRIN lens was switched from the Inscopix 9 mm GRIN lens to the GRINtech 9 mm GRIN lens.

Two-photon imaging characteristics of the GRINtrode imaging systems were measured by imaging calibration targets at an excitation wavelength of 920 nm and emission collected through a 525/50 nm bandpass filter (Semrock PN FF03-525/50-25). Magnification was characterized by imaging a 50 μm grid slide calibration target (Max Levy II-VI Aerospace and Defense, DA113) without GRIN lens imaging systems and calibrating the microns/pixel of the microscope, then imaging the 50 μm grid slide through GRIN lens imaging systems and measuring the factor by which the microns/pixel was impacted. The field of view (FOV) was measured by imaging the 50 μm grid slide (Fig. 2 A1, B1), subtracting the baseline fluorescence and counting the number of peaks from the grid slide fluorescence profile that exceeded 10% of the maximum fluorescence intensity. The lateral resolution of the GRINtrode imaging systems was characterized by imaging 0.5 μm diameter fluorescent beads (ThermoFisher G500)(Fig. 2 A2, B2) and calculating the average FWHM of Gaussian fits to the line profiles of 5 randomly sampled beads. The axial resolution of the GRINtrode imaging systems were characterized by taking a Z-stack of the 0.5 μm diameter fluorescent beads and calculating the average FWHM of Gaussian fits to the Z axis profiles of 5 randomly sampled beads. Axial FWHM was then corrected for magnification by multiplying the measured axial FWHM by a factor of 1/M^2^. The results are summarized in Fig. 2C.

**Figure 2.**
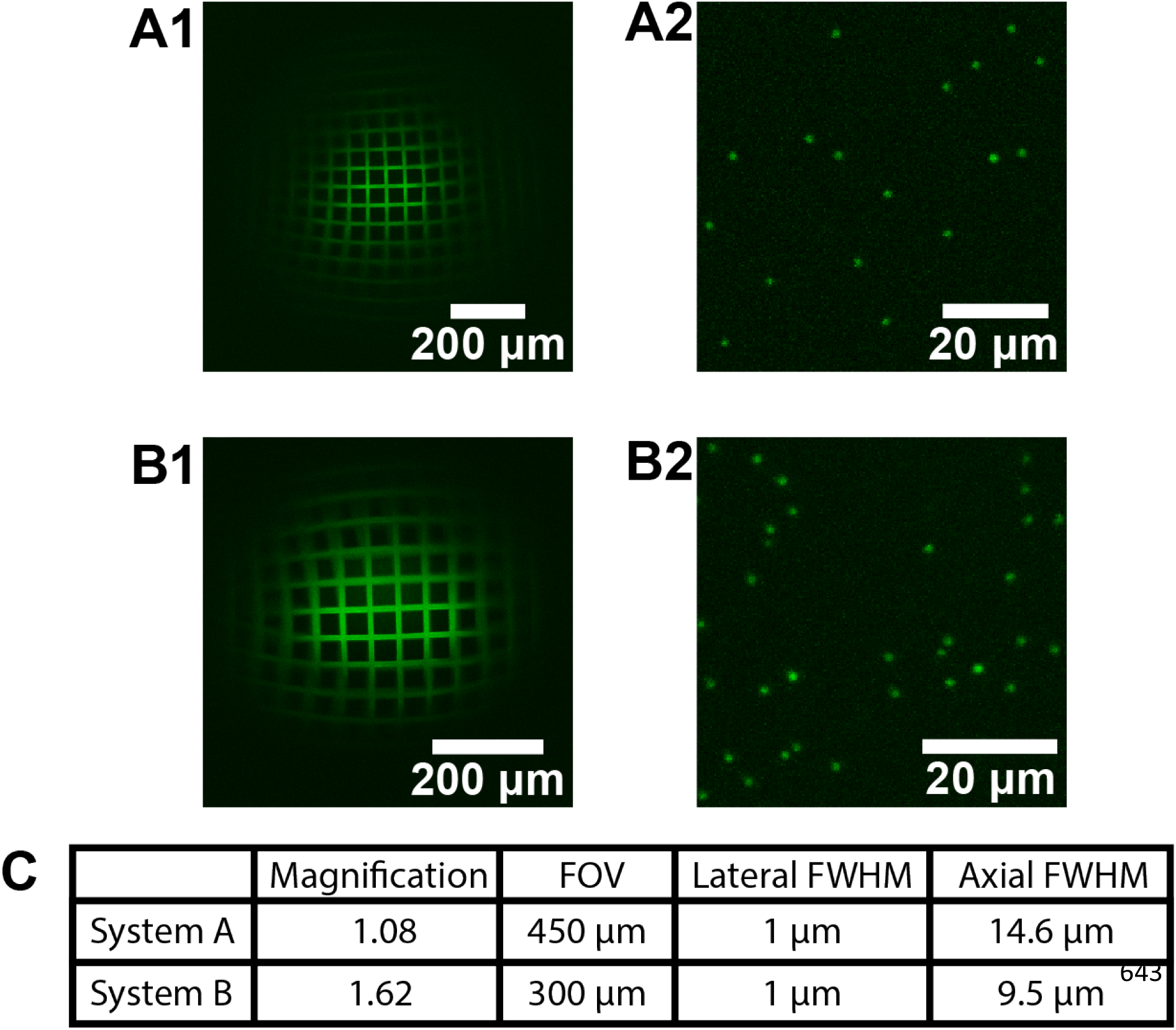
Two-photon optical characterizations of the GRINtrode imaging systems at 920 nm. Maximum intensity z-projection of a z-stack through a fluorescent 50 μm grid slide calibration target using the GRIN lens imaging system A **(A1)** and imaging system B **(B1)**. Maximum intensity z-projection a z-stack through the 0.5 μm fluorescent bead slide sample using the GRIN lens imaging system A **(A2)** and imaging system B **(B2). (C)** Table summarizing results for optical characterizations using the 50 μm grid slide for magnification and FOV, and 0.5 μm fluorescent beads for lateral and axial FWHM measurements.

### 3.3 Two-photon head-fixed GRINtrode recording

We used the GRINtrode to perform simultaneous in vivo two-photon imaging at 920 nm excitation and extracellular electrical recording. The GRINtrode was inserted into the brain at two different stereotaxic coordinates from bregma (AP: -2.4 mm, ML: +1.8 mm, DV: -1.5 mm for animal A; AP: -3.16 mm, ML: +2.5 mm, DV: -1.75 mm for animal B) and was advanced in a dorsoventral trajectory from the neocortex targeting dentate gyrus for animal A and hippocampal layer CA1 for animal B. GCaMP (pGP-AAV1.Syn.jGCaMP7f.WPRE for animal A, pAAV1.Syn.GCaMP6s.WPRE.SV40 for animal B) was virally expressed in neurons using adeno associated viruses expressing the sensor under the hSyn promoter. Additionally, GCaMP was expressed transgenically in the Thy1-GCaMP6f mice used. We used both transgenic and viral GCaMP expression to maximize chances of obtaining GCaMP signal per surgery as the goal of this work was to demonstrate feasibility of the GRINtrode device; use of a more specific GCaMP expression protocol will be crucial when addressing questions regarding unique cell-type function. Two-photon head-fixed imaging was performed using a homebuilt Sutter Instruments Movable Objective Microscope (section 2.2) and tetrode recording was performed using an Intan RHD2000 system (section 2.3). Fig. 3 shows images of GCaMP fluorescence, traces of fluorescence intensity for a subset of the regions of interest and a power spectrogram. The FOVs showed 40 and 100 areas for dataset A and B respectively with fluctuating intensity above background that were classified as different components using CaImAn (Fig. 3 A1, B1). The components had a signal to noise ratio (SNR) of 20 and above (Fig. 3 A2, B2). The wavelet power spectrogram showed increases in power that took place at low frequency as expected (Fig. 3 A3, B3).

**Figure 3.**
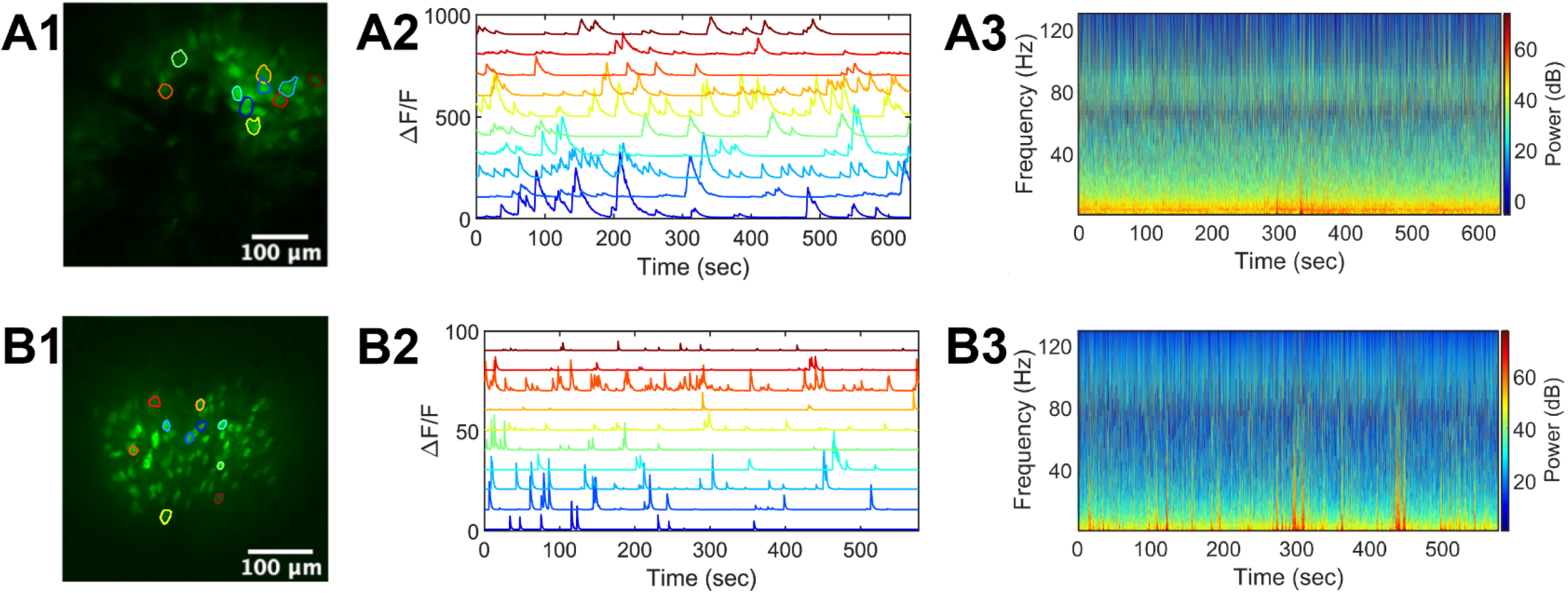
Example GRINtrode data recorded from head-fixed mice. **(A1)** Maximum intensity timelapse projection of GCaMP data from a 10-minute recording session in a female Thy1xGCaMP6f mouse with viral expression of jGCaMP7f. The GRINtrode was implanted to dentate gyrus. Selected cell ROIs are outlined. **(A2)** Color matched ΔF/F traces of GCaMP fluorescence from selected cells in A1. **(A3)** Power spectrogram of LFP activity, averaged over all 16 electrodes of the GRINtrode for the recording session. **(B1)** Maximum intensity timelapse projection of GCaMP data from a 10-minute recording session in a female Thy1xGCaMP6f mouse with viral expression of GCaMP6s. The GRINtrode was implanted to hippocampal layer CA1. Selected cell ROIs are outlined. **(B2)** Color matched ΔF/F traces of GCaMP fluorescence from selected cells in B1. **(B3)** Power spectrogram of LFP activity, averaged over all 16 electrodes of the GRINtrode for the recording session.

### 3.4 Cross-Correlation Analysis of GCaMP and LFP Data

GCaMP fluoresence of neurons imaged by the GRINtrode is expected to be directly related to action potential firing^5,16,23,24^. What would be the relationship of this optical recording of neuronal activity to the low frequency bandwidth (1-100 Hz) LFP recorded by the surrounding tetrodes? The LFP recorded by the tetrodes is generated by postsynaptic potentials in nearby neurons^4^. Interestingly, the LFP does not simply reflect the activity of nearby neurons because postsynaptic potentials that generate local current dipoles will result from the firing of nearby neurons forming local, recurrent connections as well as the firing of remote neurons with afferent inputs into a region and because there is a contribution to the LFP from distant signals through volume conduction^4,25,26^. As a result, it is expected that optical GCaMP and LFP data from the GRINtrode should be ***partially*** overlapping.

In order to evaluate overlap between optical recordings from specific cells and LFP recordings, we performed cross-correlation analysis on the data presented in Figure 3B computing the Pearson’s linear correlation coefficient between single component ΔF/F traces of GCaMP fluorescence and the LFP power traces per electrode. LFP data was split into 4 bandwidths^27^: theta (6-14 Hz), beta (15-30 Hz), low gamma (35-55 Hz) and high gamma (65-95 Hz). Cross-correlation was then computed between all 100 GCaMP traces for the timelapse, and each of these 4 bandwidths for all 16 electrodes. Figure 4C shows ΔF/F traces of the 5 optically recorded cells with the highest correlation coefficiencts to the theta bandwidth LFP data. Figures 4B and 4D show the histograms of the correlations between all pairs of LFP power and GCaMP ΔF/F traces (original histograms) compared to histograms for correlations calculated after time-shuffling of the ΔF/F traces for the theta bandwidth. Shuffling was performed 100 times by a circular shift of the trace by a random number of time points with a minimum shift of one tenth of the length of the trace. The original histograms are skewed towards positive correlations indicating that a subset of these pairwise cross-correlations are positively correlated. Indeed, when tested for significance using a KS test we find that the original histograms differ from the shuffled histograms (KS test p value <0.001). Furthermore, 75 to 90% of the correlations computed between the LFP power and dF/F traces yield p values below a significance p value adjusted using the false discovery rate to correct for multiple comparisons^28^ (Fig. 4F).

**Figure 4.**
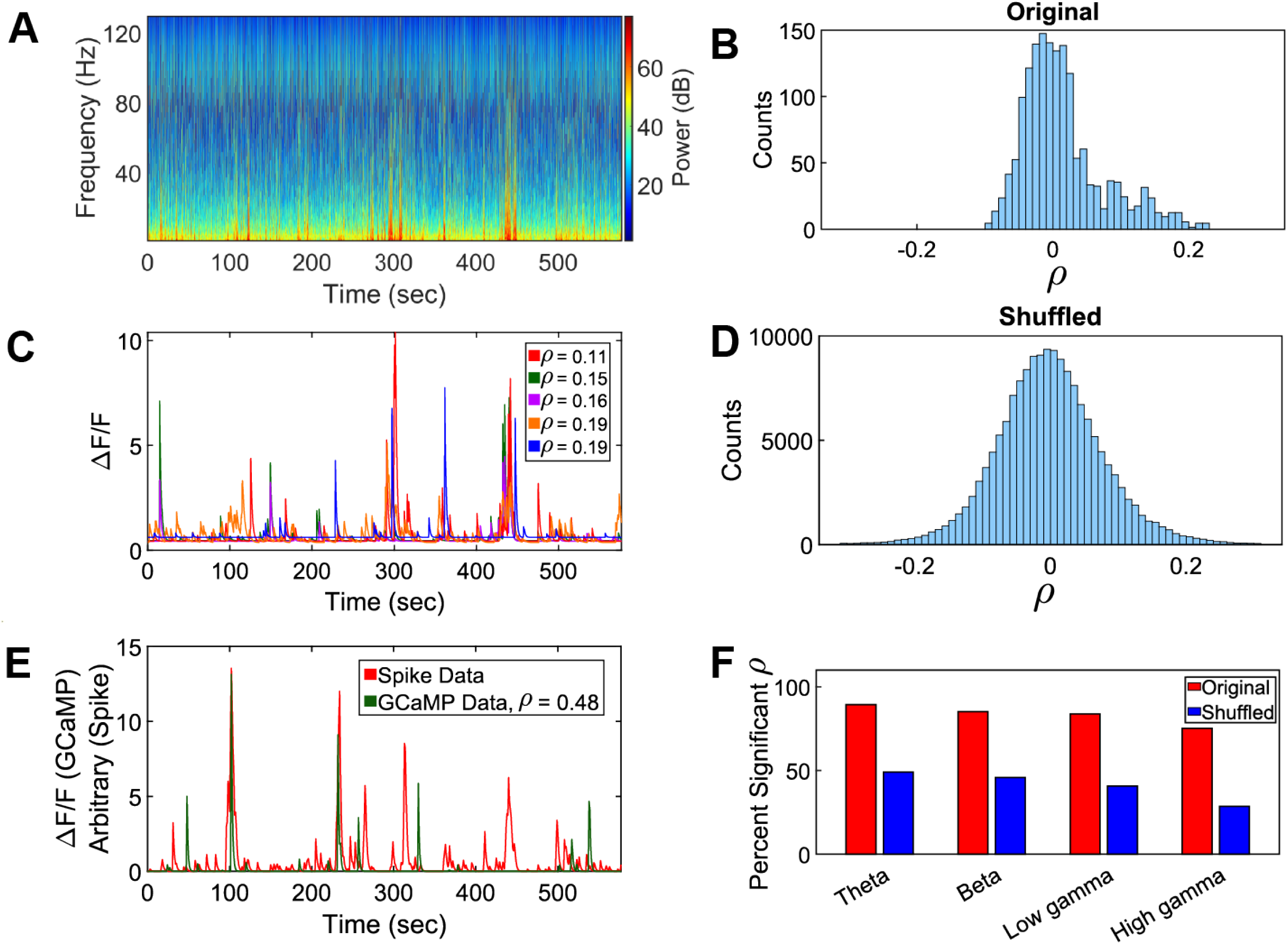
GCaMP vs. electrical data cross-correlation results analyzing data in Figure 3B. **(A)** Mean LFP power spectrogram of all 16 electrodes during a 10-minute timelapse. **(B)** Histogram of Pearson linear correlation coefficients between all 100 GCaMP cells and all 16 electrodes for the theta bandwidth, showing a skew towards positive correlations. **(C)** ΔF/F traces of the 5 optically recorded cells with highest correlation coefficients to the theta bandwidth LFP data. In the legend we are reporting the average ρ for that GCaMP trace across all 16 electrodes. **(D)** Histogram of Pearson linear correlation coefficients between shuffled GCaMP cells and all 16 electrodes for the theta bandwidth, showing a more normal distribution than the original, non-shuffled cross-correlation analysis histogram. **(E)** Cross-correlation analysis results between GCaMP data and convolved spike data using Figure 3C data. Binary spike data was convolved with an exponential decay function using the same decay constant used in CaImAn analysis to approximate what the activity of these spiking cells would look like as GCaMP signal. We then used Pearson’s linear correlation coefficient to find the optically recorded cells which match the processed spike data most closely. Here we are showing the result with the highest correlation coefficient, ρ = 0.475. The units of the spike data are arbitrary, so we have scaled the spike data trace to be of similar amplitude to the GCaMP trace. **(F)** Bar chart showing per bandwidth percentage of significant correlations for Pearson correlation coefficient analysis for original and shuffled data sets.

**Figure 5.**
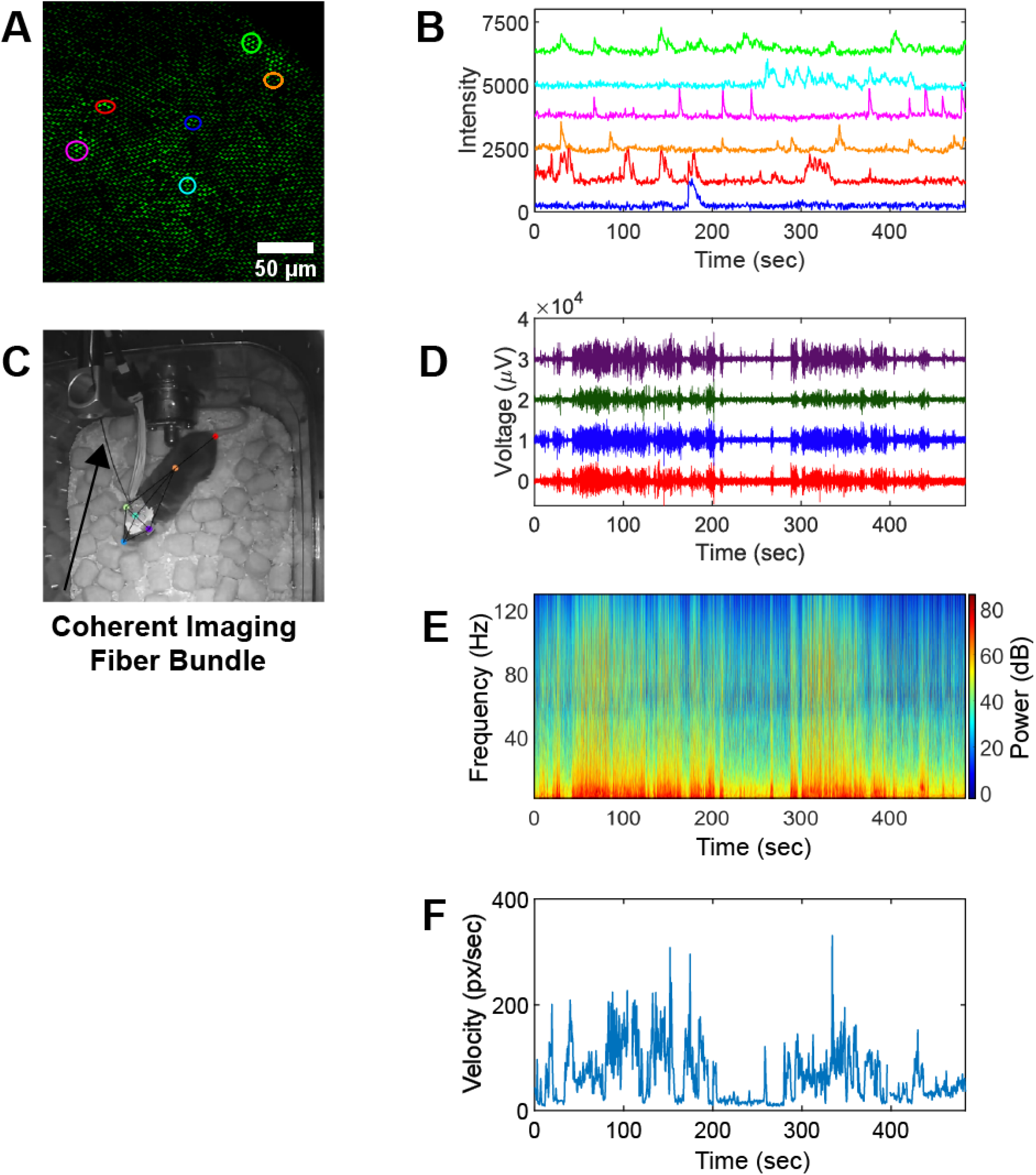
Fiber coupled freely moving two-photon GRINtrode recording. **(A)** Standard deviation timelapse projection of an 8-minute recording session, highlighted ROIs are color coordinated to traces in B. **(B)** Color matched fluorescence intensity traces from ROIs in A, time synced to D, E and F. **(C)** Still frame of behavioral video showing freely moving fiber coupled animal and markers generated by analysis with DeepLabCut. **(D)** Raw LFP activity from the 4 tetrodes of the GRINtrode, time synced to B, E and F. **(E)** Mean LFP power spectrogram of all 16 electrodes, time synced to B, D and F. **(F)** Velocity of the head of the mouse during the behavioral video generated by DeepLabCut analysis, time synced to B, D and E.

#### 3.4.1 Cross-Correlation Analysis of GCaMP and Spike Data

Since spiking activity from multiple neurons neighboring an electrode contribute to the high frequency (500-5000Hz) filtered field potential^2,3,4^ it may theoretically be possible to record the same neuron both optically and electrically using the GRINtrode if the electrical and optical recording fields are overlapping. There are challenges in achieving overlapping optical and electrical fields though. The recording plane of the tetrodes relative to the imaging plane is determined by the length of tetrode that extends pass the implanted GRIN lens. We attempt to make this length as close to the working distance of the GRINtrode imaging system as possible, however since tetrodes are cut by hand there is variability in this parameter. Additionally, tetrodes are fixed on the periphery of the GRIN lens, 500 μm laterally from the center of the imaging field and provide a theoretical recording sphere radius of 100 μm^29^. With an FOV of ∼450 μm, overlap between optical and electrical recording fields is not expected. Still, it may be the case that optical and electrical single units display correlated activity even though they come from different recording fields. With these caveats in mind, we proceeded to explore the relationship between single unit electrical activity and GCaMP fluorescence changes. We extracted single unit activity from electrical recordings using a custom implementation^30^ of the wave_clus software package^18^. The software outputs a binary timeseries, with single unit spike timings represented as a one. After extracting spikes, we then convolved this binary series with an exponential decay function using the same decay constant used in the CaImAn^16^ analysis of the optical recordings to approximate what the spike activity would look like converted to a GCaMP signal. We then performed cross-correlation analysis using Pearson’s linear correlation coefficient to seek out optically recorded cells that match closest to the processed spiking activity. Fig. 4E shows that the convolved firing of one isolated single unit displays a Pearson correlation coefficient of 0.48. The spike and GCaMP data show two temporally localized transients, but several spike transients that are not accounted for by the GCaMP data. We suspect this is because the isolated “single unit” was not in the imaging plane and displayed correlated spiking with units in the imaging plane, perhaps connected through a neuronal network.

### 3.5 Freely Moving Two-Photon Fiber Coupled GRINtrode Recording

We performed freely moving two-photon imaging using a 1.5 meter long Fujikura FIGH-15-600N coherent imaging fiber bundle inserted into the imaging optic channel^15^. A single mode fiber and grating-pair compressor were used to correct for the linear and nonlinear dispersion of fiber bundle (see Methods), allowing short pulses at the sample. This set up allowed us to perform simultaneous two-photon imaging of GCaMP activity and extracellular electrophysiology with the GRINtrode in a freely moving mouse. The GRINtrode was inserted into the brain at stereotaxic coordinates (AP: -2.4 mm, ML: +1.8 mm, DV: -1.25 mm) referenced to bregma in a 3-month-old Thy1-GCaMP6f mouse and was advanced in a dorsoventral trajectory from the neocortex targeting the CA1 layer of the hippocampus. The calcium sensor jGCaMP7f was expressed in neurons using AAV under the hSyn promoter. Two-photon fiber coupled freely moving imaging was performed using a custom-built microscope (section 2.2) and tetrode recording was performed using an Intan RHD2000 system (section 2.3). A Sutter MPC-325 micromanipulator system was used to fine adjust the Z spacing between the chronically implanted GRIN lens inside of the GRINtrode and the coherent imaging fiber bundle, and the fiber bundle was locked in place using the imaging optic set screws.

The mouse movement was recorded with a camera, while simultaneous acquiring two-photon imaging (Fig. 5A, 5B) and LFP recording (Fig. 5D, 5E) as the mouse foraged through a cage with food pellets (Fig. 5C, Movie 1). We extracted behavioral kinematics using the DeepLabCut software package^19^. Figs. 5B, 5D and 5E show temporally synchronized GCaMP fluorescence intensity traces, raw LFP traces, and LFP spectrogram while the mouse foraged through the cage. Fig. 5F shows the head velocity of the mouse during the recording session. Together, these results demonstrate the utility of the GRINtrode in providing simultaneous two-photon imaging and extracellular electrophysiology in a freely moving mouse.

## 4 Discussion

We developed and validated a GRINtrode, the first GRIN lens and tetrode based head-mounted neural implant designed for simultaneous two-photon calcium imaging and extracellular electrophysiology. Importantly, we have demonstrated the feasibility of fiber-coupling the GRINtrode to allow imaging and electrical recording of neural activity in the freely moving mouse. Simultaneously recording neural activity with optical and electrical modalities provides a unique opportunity to probe the relationship between these recording methods in vivo. We have demonstrated positive correlation between optically and electrically recorded activity by performing cross-correlation analyses between the two data types.

### 4.1 Extracellular Field Potential Electrophysiology

Extracellular field potential recordings are the oldest method of recording in vivo neural activity, and as such there is an abundance of knowledge based on this technique. Therefore, it is desirable for neurophysiologists to collect optical data with simultaneous electrophysiology in vivo to build upon prior knowledge obtained in systems neuroscience. By placing electrodes into the brain tissue, researchers can record a direct measure of neural activity relayed in the form of action potentials. Action potentials create current sinks and sources whose flow through the resistive extracellular compartment is detected as voltage changes at the recording site. The recording site of single electrodes can be used to capture neural oscillations, a signature of coordinated dynamics in the brain^4,31^ generated by the activity of large ensembles of neurons. While such information is crucial to decoding the underlying rhythms of the brain, such recordings have a trade-off of high temporal resolution and poor spatial resolution^32^. In coupling with optical methods, researchers can simultaneously explore the relationship between single cell responses and network ensembles, revealing new insights into brain-state dynamics at an intermediary scale.

### 4.2 Optical Neural Activity Recording

Optical methods of recording in vivo neural activity are a relatively new method in the field of neuroscience that have become popular because they allow recording from large ensembles of spatially resolved neurons^11,33–36^. Optically recording neural activity requires a fluorescent reporter whose fluorescence intensity is modulated by neural activity. There are several classes of these reporters: calcium indicators, neurotransmitter indicators and voltage indicators. In this work, we focus on the use of the widely used calcium indicator, GCaMP^24^.

Optical recording methods are subject to some limitations due to the scattering nature of brain tissue, restricting imaging depth. However, there are methods to overcome these limitations. Multiphoton imaging uses longer wavelength excitation light, such that multiple coincident photons are needed to excite the fluorophore. The use of longer near-infrared wavelengths increases optical penetration due to reduced scattering compared with visible wavelengths. Multiphoton imaging also provides optical sectioning, as the probability of fluorophore excitation depends on intensity which is confined to the focus of the imaging system.

Additionally, scattering tissue can be bypassed by implanting imaging optics in deep brain regions. Gradient Refractive Index (GRIN) lenses are rod shaped lenses which provide optical power by means of a radially dependent index of refraction. These lenses are useful for in vivo neural imaging in deeper brain regions that are otherwise not optically accessible due to the limited penetration depth of optical imaging. Their small form factor allows them to be chronically implanted above the region of interest^21^. One significant trade-off of this approach is the invasiveness of implantation and the surgical skill needed to capture optical responses in vivo. GRIN lens implantation requires traumatic surgeries in rodents, where the surgeon aspirates superficial structures above the region of interest prior to GRIN lens insertion.

However, when using proper surgical techniques during implantation, the datasets collected with the GRINtrode are rich and can convey unseen data compared to use of a single neural recording modality alone. Therefore, the larger need to record electrical and optical activity at cellular and network resolution in deep brain regions can be performed using the GRINtrode, yet its insertion with minimal disturbance to brain structures superficial to the recording site remains impossible for now.

Optical techniques often require head-fixing the animal to perform imaging, limiting the behavioral paradigms capable of being investigated. If the entire imaging system can be miniaturized to the extent that the animal can carry it, freely moving wireless imaging is possible^37^. This is currently limited to 1 photon imaging modalities though, as sufficiently miniaturized coherent laser sources are not available. However, it is possible to use optical fibers to fiber couple imaging systems and allow for multiphoton imaging in freely moving animals^15^.

#### 4.3 Bringing together the information encoded by LFP recording and optical imaging of neuronal activity

Extracellular LFP recording has provided important information on the behavioral relevance and circuit mechanisms of neural function^4,27,38,39^. High frequency bandwidth recordings (500-5000 Hz) yield information on single neuron activity particularly when recording is performed with multiunit electrodes making it more convenient to isolate single units^14,40^. Furthermore, lower frequency bandwidth LFP (1-250 Hz) also provides key information on neural circuits although understanding of the relationship between the LFP and the activity of single neurons is difficult because of a lack of a well-posed inverse model to determine single unit activity based on conservation of charge and Maxwell’s equations^4^. Development of miniature devices that allow simultaneous imaging of neural activity and electrical recording of the LFP provides the new tools to understand the relationship between the LFP and single neuron activity and provides the opportunity to use closed loop modulation based on either the LFP or on single unit activity and find the consequence on neural activity and behavior^38,39,41^.

Our GRINtrode device provides the opportunity for simultaneous optical imaging and field potential recording in the freely moving animal. Tetrodes are bundles of 4 electrodes which provide multiple recording sites with a ∼20 μm spatial separation. These multiple recording sites allow for isolation of single unit spiking activity as spike amplitude is a function of distance to the recording site. However, use of tetrodes limits the number of single units that can be isolated and the ability to achieve overlapping optical and electrical fields. Future versions can incorporate patterned multielectrode devices to provide extraction of more single units and accurate control over relative placement of optical and electrical recording fields.

## 5 Conclusion

We demonstrate a robust, easy to construct, and novel neural implant, the GRINtrode, that allows for simultaneous optical and electrical recording of neuronal activity in vivo. It is suitable for head-fixed one-photon or two-photon optical recording with simultaneous extracellular electrophysiology provided by 16 electrodes grouped into 4 tetrodes. We also demonstrated two-photon fiber-coupled GRINtrode recording, allowing use in freely moving animal recording.

There has been increasing effort to develop methods for simultaneous optical and electrophysiological recording of neural activity in vivo. There is abundant motivation to develop and improve methods for performing this recording: investigating GCaMP dynamics versus direct electrical recordings of single unit activity, using electrical recording to compensate for the limits on temporal resolution intrinsic to optical methods, development of closed loop optogenetic systems based on LFP activity. The GRINtrode presents a versatile solution to this desire for bimodal recording, it is easy to construct and uses a commonplace method for electrical recording and allows for advanced optical techniques such as multiphoton imaging and freely moving recording by means of fiber coupling capability.

## Acknowledgments

This research was supported by an Administrative Supplement to NIH UF1 NS116241, NIH R01 DC000566, and NSF BCS-1926676.

Polyimide tubing was courteously provided by MicroLumen Inc.

## Code, Data, and Materials Availability

The authors have made code used for analysis presented in this manuscript available for access through GitHub/restrepd/drta, GitHub/restrepd/drgMaster and GitHub/ConMark/GRINtrode/Code. Data presented in this manuscript is available for access through author request. CAD models of GRINtrode components are available for access through GitHub/ConMark/GRINtrode/CAD.

## References

1. Adrian, E. D. The electrical activity of the mammalian olfactory bulb. Electroencephalogr. Clin. Neurophysiol. 2, 377–388 (1950).

2. Bédard, C., Kröger, H. & Destexhe, A. Modeling Extracellular Field Potentials and the Frequency-Filtering Properties of Extracellular Space. Biophys. J. 86, 1829–1842 (2004).

3. Buzsáki, G., Anastassiou, C. A. & Koch, C. The origin of extracellular fields and currents — EEG, ECoG, LFP and spikes. Nat. Rev. Neurosci. 2012 136 13, 407–420 (2012).

4. Pesaran, B. et al. Investigating large-scale brain dynamics using field potential recordings: analysis and interpretation. Nat. Neurosci. 2018 217 21, 903–919 (2018).

5. Chen, T. W. et al. Ultrasensitive fluorescent proteins for imaging neuronal activity. Nat. 2013 4997458 499, 295–300 (2013).

6. Wu, Z., Lin, D. & Li, Y. Pushing the frontiers: tools for monitoring neurotransmitters and neuromodulators. Nat. Rev. Neurosci. 2022 235 23, 257–274 (2022).

7. Knöpfel, T. & Song, C. Optical voltage imaging in neurons: moving from technology development to practical tool. Nat. Rev. Neurosci. 2019 2012 20, 719–727 (2019).

8. Platisa, J. & Pieribone, V. A. Genetically encoded fluorescent voltage indicators: are we there yet? Curr. Opin. Neurobiol. 50, 146 (2018).

9. Salinas, E. & Sejnowski, T. J. Correlated neuronal activity and the flow of neural information. Nat. Rev. Neurosci. 2001 28 2, 539–550 (2001).

10. Thunemann, M. et al. Deep 2-photon imaging and artifact-free optogenetics through transparent graphene microelectrode arrays. Nat. Commun. 9, 1–12 (2018).

11. Aharoni, D., Khakh, B. S., Silva, A. J. & Golshani, P. All the light that we can see: a new era in miniaturized microscopy. Nat. Methods 2018 161 16, 11–13 (2018).

12. Wu, X. et al. A Modified Miniscope System for Simultaneous Electrophysiology and Calcium Imaging in vivo. Front. Integr. Neurosci. 0, 21 (2021).

13. Cobar, L. F., Kashef, A., Bose, K. & Tashiro, A. Opto-electrical bimodal recording of neural activity in awake head-restrained mice. Sci. Reports 2022 121 12, 1–16 (2022).

14. Gray, C. M., Maldonado, P. E., Wilson, M. & McNaughton, B. Tetrodes markedly improve the reliability and yield of multiple single-unit isolation from multi-unit recordings in cat striate cortex. J. Neurosci. Methods 63, 43–54 (1995).

15. Ozbay, B. N. et al. Three dimensional two-photon brain imaging in freely moving mice using a miniature fiber coupled microscope with active axial-scanning. Sci. Rep. 8, (2018).

16. Giovannucci, A. et al. CaImAn an open source tool for scalable calcium imaging data analysis. Elife 8, 1–45 (2019).

17. Schindelin, J. et al. Fiji: an open-source platform for biological-image analysis. Nat.Methods 2012 97 9, 676–682 (2012).

18. Quiroga, R. Q., Nadasdy, Z. & Ben-Shaul, Y. Unsupervised spike detection and sorting with wavelets and superparamagnetic clustering. Neural Comput. 16, 1661–1687 (2004).

19. Mathis, A. et al. DeepLabCut: markerless pose estimation of user-defined body parts with deep learning. Nat. Neurosci. 21, 1281–1289 (2018).

20. Dana, H. et al. Thy1-GCaMP6 transgenic mice for neuronal population imaging in vivo. PLoS One 9, (2014).

21. Resendez, S. L. et al. Visualization of cortical, subcortical and deep brain neural circuit dynamics during naturalistic mammalian behavior with head-mounted microscopes and chronically implanted lenses. Nat. Protoc. 11, 566–597 (2016).

22. Anikeeva, P. et al. Optetrode: A multichannel readout for optogenetic control in freely moving mice. Nat. Neurosci. 15, 163–170 (2012).

23. Meng, G. et al. High-throughput synapse-resolving two-photon fluorescence microendoscopy for deep-brain volumetric imaging in vivo. Elife 8, 1–24 (2019).

24. Dana, H. et al. High-performance calcium sensors for imaging activity in neuronal populations and microcompartments. Nat. Methods 16, 649–657 (2019).

25. Kajikawa, Y. & Schroeder, C. E. How local is the local field potential? Neuron 72, 847–858 (2011).

26. Herreras, O. Local field potentials: Myths and misunderstandings. Front. Neural Circuits 10, 101 (2016).

27. Losacco, J., Ramirez-Gordillo, D., Gilmer, J. & Restrepo, D. Learning improves decoding of odor identity with phase-referenced oscillations in the olfactory bulb. Elife 9, (2020).

28. Curran-Everett, D. Multiple comparisons: Philosophies and illustrations. Am. J. Physiol. - Regul. Integr. Comp. Physiol. 279, (2000).

29. Jog, M. S. et al. Tetrode technology: advances in implantable hardware, neuroimaging, and data analysis techniques. J. Neurosci. Methods 117, 141–152 (2002).

30. Li, A., Gire, D. H. & Restrepo, D. ϒ Spike-Field Coherence in a Population of Olfactory Bulb Neurons Differentiates between Odors Irrespective of Associated Outcome. J. Neurosci. 35, 5808–5822 (2015).

31. Friston, K. J., Bastos, A. M., Pinotsis, D. & Litvak, V. LFP and oscillations-what do they tell us? Current Opinion in Neurobiology vol. 31 1–6 (2015).

32. Swanson, J. L. et al. Advancements in the quest to map, monitor, and manipulate neural circuitry. Front. Neural Circuits 0, 45 (2022).

33. Zong, W. et al. Large-scale two-photon calcium imaging in freely moving mice. Cell 185, 1240–1256.e30 (2022).

34. Yu, C. H., Stirman, J. N., Yu, Y., Hira, R. & Smith, S. L. Diesel2p mesoscope with dual independent scan engines for flexible capture of dynamics in distributed neural circuitry. Nat. Commun. 2021 121 12, 1–8 (2021).

35. Helmchen, F. & Denk, W. Deep tissue two-photon microscopy. Nat. Methods 2005 212 2, 932–940 (2005).

36. Sofroniew, N. J., Flickinger, D., King, J. & Svoboda, K. A large field of view two-photon mesoscope with subcellular resolution for in vivo imaging. Elife 5, (2016).

37. Barbera, G., Liang, B., Zhang, L., Li, Y. & Lin, D. T. A wireless miniScope for deep brain imaging in freely moving mice. J. Neurosci. Methods 323, 56–60 (2019).

38. Siegle, J. H. & Wilson, M. A. Enhancement of encoding and retrieval functions through theta phase-specific manipulation of hippocampus. Elife 2014, (2014).

39. Kanta, V., Pare, D. & Headley, D. B. Closed-loop control of gamma oscillations in the amygdala demonstrates their role in spatial memory consolidation. Nat. Commun. 2019 101 10, 1–14 (2019).

40. Steinmetz, N. A. et al. Neuropixels 2.0: A miniaturized high-density probe for stable, long-term brain recordings. Science (80-.). 372, (2021).

41. Grosenick, L., Marshel, J. H. & Deisseroth, K. Closed-Loop and Activity-Guided Optogenetic Control. Neuron 86, 106 (2015).

