## Supplemental for "GRINtrode: A neural implant for simultaneous two-photon imaging and extracellular electrophysiology in freely moving animals"

*S1 One-Photon Fiber-coupled GRINtrode recording*

Head-fixing an animal induces a stress response and severely limits the behavioral tasks an animal can perform. It is desirable to allow animals to be freely moving to avoid this stress response and allow for a broader selection of behavioral tasks. To address these concerns the GRINtrode was designed to allow for coupling to a coherent fiber imaging bundle to allow for freely moving optical recording.

To demonstrate the feasibility of freely moving recording and the fiber coupling capabilities of the GRINtrode, we inserted a doublet GRIN lens (GRINtech NEM-100-25-10-860-DS) coupled to a coherent fiber imaging bundle (Fujikura FIGH-15-600N) into the imaging optic channel of the GRINtrode in a head-fixed animal and used a Nikon A1R confocal microscope to perform imaging. An Olympus UPlanXApo 10X objective (Olympus, PN N5701900) is used to focus the imaging laser onto the cores of the fiber-bundle, which is housed in a XYZ translator (Thorlabs CXYZ1) to allow for adjustment of the fiber position. As this was a proof-of-concept demonstration of one-photon fiber coupled imaging, we did not record tetrode data for the experiment.

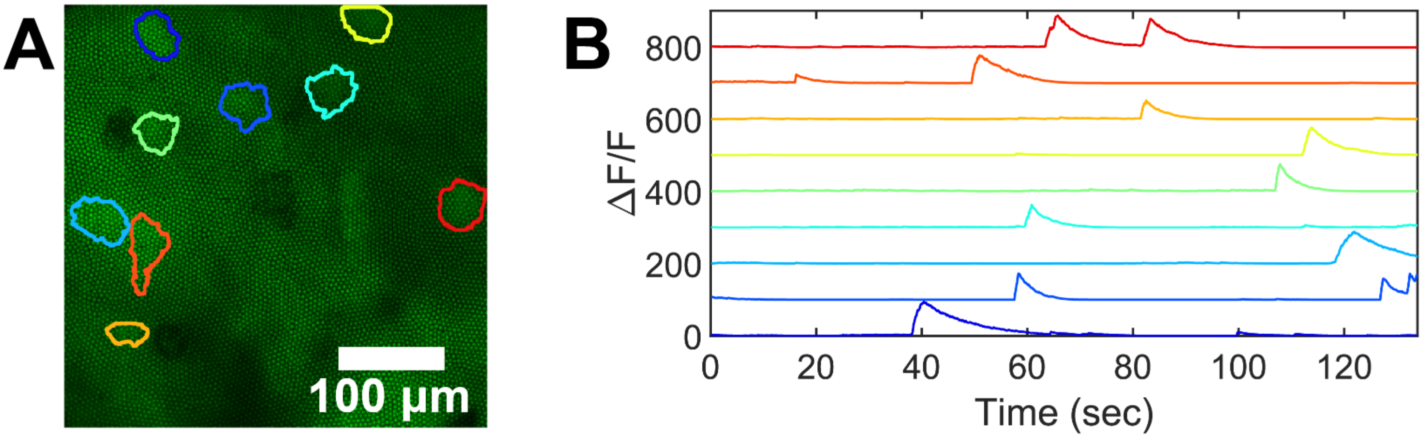


**Figure S1.** 1-Photon fiber-coupled GRINtrode data. **(A)** Maximum intensity projection of a 2-minute timelapse recorded using the fiber-coupled GRINtrode targeting dentate gyrus in a female Thy1xGCaMP6f mouse with viral expression of jGCaMP7f. Selected cell ROIs are outlined. **(B)** Color matched ΔF/F traces of GCaMP fluorescence from cell ROIs in A.

*S2 FOV and Resolution Characterizations*

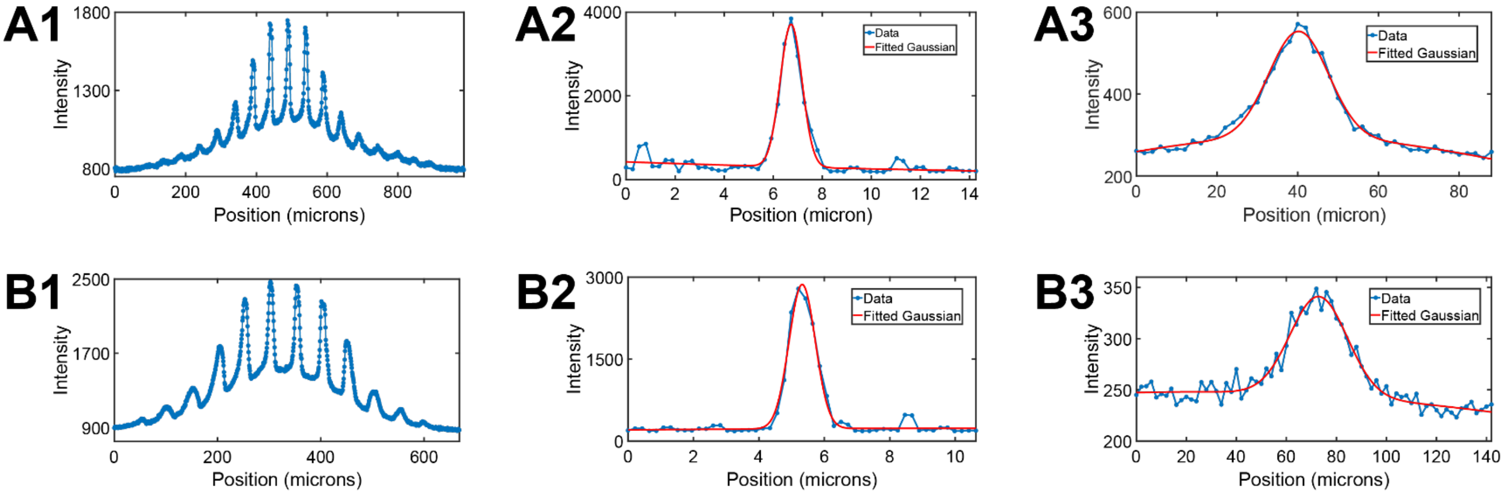


**Figure S2.** FOV and lateral/axial FWHM measurements. **(A1)** Field profile of 50 µm pitch fluorescent grid target using GRIN lens imaging system A. **(A2)** Example Gaussian fit for the lateral fluorescence intensity profile of one 0.5 µm fluorescent bead using system A, average FWHM calculated over 5 randomly selected 0.5 µm fluorescent beads = 1 µm. **(A3)** Example Gaussian fit for the axial fluorescence intensity profile of one 0.5 µm fluorescent bead using system A, average axial FWHM calculated over 5 randomly selected 0.5 m fluorescent beads = 17 µm. After correcting for magnification, FWHM = 14.6 µm**. (B1)** Field profile of 50 µm pitch fluorescent grid target using GRIN lens imaging system B. **(B2)** Example Gaussian fit for the lateral fluorescence intensity profile of one 0.5 µm fluorescent bead using system B, average FWHM calculated over 5 randomly selected 0.5 µm fluorescent beads = 1 µm. **(B3)** Example Gaussian fit for the axial fluorescence intensity profile of one 0.5 µm fluorescent bead using system B, average axial FWHM calculated over 5 randomly selected 0.5 µm fluorescent beads = 25 µm. After correcting for magnification, FWHM = 9.5 µm**.**

**GRINtrode Assembly Protocol**

Supplies:

- GRINtrode body
- 4 tetrodes (prepared using Sandvik Precision Fine Tetrode Wire, Sandvik PN PX000001; built using the method presented in Gray et al. (1995)[14].
- Neuralynx EIB-16 board (https://neuralynx.com/hardware/eib-16)
- Large EIB gold pins (https://neuralynx.com/hardware/large-eib-pins)
- Tetrode pinning tool (fine forceps or insect pin)
- Ground wire (https://www.fishersci.com/shop/products/bare-silver-wire-0-008/NC0326296)
- Implanted GRIN lens (typically 9 mm x 1mm Inscopix or GRINtech, can change to a doublet if magnification is desired)
- MicroLumen 400-II PI tubing (ID=0.401", OD=0.0413")
- MicroLumen 450-II PI tubing (ID=0.0451", OD=0.0490")
- MicroLumen 575-I PI tubing (ID=0.0577", OD=0.0598")
- Fine forceps
- Vannas scissors
- Bulldog forceps with heat shrink on tips (for holding GRIN lenses)
- Loctite super glue ultra-gel minis (high viscosity super glue) (https://www.amazon.com/Loctite-Super-Glue-Ultra-Minis/dp/B01FIL87AK?th=1)
- Metabond (Clear/Radiopaque L-powder, catalyst, quick base, ceramic dish) (https://www.parkell.com/C-B-Metabond-Quick-Adhesive-Cement-System)
- Two-part epoxy

(https://www.amazon.com/Gorilla-Epoxy-Minute-ounce-Syringe/dp/B001Z3C3AG)

- Kwil-Sil Silicone Elastomer (WPI Inc, KWIK-SIL) (https://www.wpiinc.com/kwik-sil-low-toxicity-silicone-adhesive)
- Parafilm
- Lens tissue
- Methanol
- Acetone
- Quadhands or similar helping hands holder tool (https://www.quadhands.com/products/quadhands)
- Digital calipers
- #52 drill bit
- Insect pins
- GRIN lens holding plate
- Dissecting scope
- Cotton tipped applicators

Procedure:

1. Inspect GRIN lenses and clean with methanol or acetone (acetone for metabond or epoxy removal if needed, can soak)
2. Cut 1 piece of 400-II tubing to ½ - ¾ the length of your implanted GRIN lens
3. Using insect pins, mix and place very small dot of 2 part epoxy around your implanted GRIN lens. Using bulldog forceps with heatshrink over lens paper, push the 400-II tubing over the lens to be slightly past flush with the end of the GRIN lens. Let dry.
4. Remove the dowel pins from the GRINtrode body and take the thumb screw and vented screw out of the GRINtrode body. Work with the thumb screw and vented screw until reassembly is mentioned.

5. Cut 1 piece of 575-I tubing to total height of device with screw and most retracted point (around 11.5-12 mm) + desired implant depth + 0.75 mm for EIB board
    a. For safer epoxy application, set the thumb screw so that the vented screw is just engaging the GRINtrode walls. Remove the thumb screw and insert the tubing into the bottom of the vented screw, lightly applying epoxy and removing excess as you go, until you have around 1 mm sticking out of the top of the vented screw. Cut the tubing with around (implant depth + 3) mm sticking out of bottom of the vented screw. Let dry.
6. Cut one piece of 450-II tubing to (implant depth + 10-15) mm
7. Using bulldog forceps to hold the bare glass end of the GRIN lens and 400-II tube assembly, slip the 450-II tubing over the end with 400-II tubing. Using insect pins lightly apply epoxy to the 400-II tube and slide the 450-II tube until only 0.5 – 1 mm of the 400-II tubing is uncovered. Let dry.
8. Reassemble the GRINtrode by placing the thumb screw & vented screw into the GRINtrode body and reinserting the friction dowels.
9. Set the digital calipers to your desired implant depth and place the depth rod on the bottom of the GRINtrode body. Using the vannas scissors slowly cut down the 575-I tubing sticking out of the bottom of the vented screw until it matches your implant depth (height of the depth rod).
10. Using a #52 drill bit, drill out one of the corner holes of the EIB-16 board to be slightly larger.
11. Hold the GRINtrode right side up in the quad hands. Lower the EIB-16 board onto the vented screw, passing the 575-I tubing through the larger corner hole of the EIB-16 board. Lower the EIB-16 board so it sits flush with the vented screw. Carefully apply metabond to adhere the EIB board to the vented screw. It is advisable to place the EIB

board diagonally facing backwards to limit animal access. Let dry and apply between the bottom of the EIB board and cap of the vented screw as well.

12. After drying, cut the 575-I tubing sticking out of the top of the EIB board to be flush.
13. Add the ground wire to the EIB-16 board by inserting the wire into one of the ground pin holes and soldering in place.
14. Insert the GRIN lens tubing assembly through the EIB board and GRINtrode so that the GRIN lens is flush with the tubing at the bottom of the device. Use the sharpie to mark the tubing at the top of this device. This serves as a reference for where to glue your tetrode relative to the loop, the start of the loop should go 1-3 mm below this depending on the size of the loop (how much spare electrode wire you'll have to get to the pin holes)
15. Under the dissection scope, use insect pins and Loctite ultra gel minis (high viscosity super glue) to adhere tetrodes. A GRIN lens holding plate will be helpful here. Begin by gluing the loop into place under the mark on the tubing, and then applying a small dot of glue on the end of the GRIN lens. Drag the tetrode into the glue on the end of the GRIN lens using forceps. Use insect pins to spread the glue around the tetrode. This helps keep the tetrode attached but also keeps the glue thin, it is vital to keep your glue layers here as thin as possible. Glue one tetrode into place, dry, rotate 180 degrees, dry, rotate 90 degrees, dry, rotate 180 degrees, dry.
16. Once tetrodes are adhered to the GRIN lens and tubing assembly, they can be cut to a more manageable length. Do not make the final cut though.
17. Holding the GRINtrode body in the quad hands, push the GRIN lens and tetrode assembly into the 575-I tubing from the top of the EIB board. If the fit is too tight, try removing excess super glue that was used on the tetrodes with a cotton tipped applicator soaked in acetone.

18. Begin lightly applying 2-part epoxy around the outside of the GRIN lens and tetrode assembly once it is ~5 mm into the device.
19. Continue lightly applying epoxy and pushing the GRIN lens and tetrode assembly down until you can barely see the GRIN lens sticking out of the bottom of the 575-I tubing at the bottom of the drive which was set to you implant depth in step 9.
20. Allow the GRIN lens and tetrode assembly to dry inside of the 575-I tubing
21. To pin the electrodes, cut a tetrode loop and isolate a single electrode. Draw the electrode into a pin hole using forceps, lightly place the gold pin, pull excess line through the pin hole, then pin the electrode using forceps or an insect pin to apply pressure to the gold pin (very little pressure is needed). Repeat until all 16 electrodes are pinned. (A 100% success rate for pinned lines is rare, 12 and above is acceptable and still worth implanting).
22. After pinning all the tetrodes, lightly apply 2-part epoxy over the top of the EIB board to protect the electrode lines. Be careful not to cover the polyimide tube or the plug of the head stage.
23. Under the dissection scope, cut the tetrodes to the desired length at the bottom of the GRIN lens.
24. Cut a ~ 1mm x 1mm square of parafilm and cover the imaging optic channel with it.
25. Use Kwik-Sil to cover the parafilm and seal the imaging optic channel.
